# Focus of attention modulates the heartbeat evoked potential

**DOI:** 10.1101/384305

**Authors:** Frederike H. Petzschner, Lilian A. Weber, Katharina V. Wellstein, Gina Paolini, Cao Tri Do, Klaas E. Stephan

**Author notes:** joint first authorship (Frederike H. Petzschner, Lilian A. Weber).

## Abstract

Theoretical frameworks such as predictive coding suggest that the perception of the body and world – interoception and exteroception – involve intertwined processes of inference, learning, and prediction. In this framework, attention is thought to gate the influence of sensory information on perception. In contrast to exteroception, there is limited evidence for purely attentional effects on interoception. Here, we empirically tested if attentional focus modulates cortical processing of single heartbeats, using a newly-developed experimental paradigm to probe purely attentional differences between exteroceptive and interoceptive conditions in the heartbeat evoked potential (HEP). We found that the HEP is significantly higher during interoceptive compared to exteroceptive attention, in a time window of 520-580ms after the R-peak. Furthermore, this effect predicted self-report measures of autonomic system reactivity. This study thus provides direct evidence that the HEP is modulated by attention and supports recent interpretations of the HEP as a neural correlate of interoceptive prediction errors.

## Introduction

Perception corresponds to inferring the causes of our sensations – whether they originate from the external world (*exteroception*) or our own bodies (*interoception*). Interoception and exteroception have long been studied in isolation. In some sense, this was natural given that they are characterized by different sensory receptors – e.g., chemo-, baro-, or thermoreceptors for interoception versus photo-, mechano-, or olfactory receptors for exteroception – and utilize distinct neural pathways for processing this information (Craig, 2002, 2009). More recently, however, theories from computational neuroscience have integrated interoception and exteroception conceptually, emphasizing common algorithmic principles and describing them as intertwined processes of inference, learning, and prediction according to probability theory or, simply speaking, Bayes’ theorem (Friston, 2009; Petzschner, Weber, Gard, & Stephan, 2017; Seth, 2013; Seth, Suzuki, & Critchley, 2012; Stephan et al., 2016).

These conceptions provide a foundation for efforts towards a more mechanistic understanding of brain-body interactions (Henningsen et al., 2018; Khalsa et al., 2018; Owens, Friston, Low, Mathias, & Critchley, 2018; Petzschner et al., 2017; Seth & Friston, 2016; Smith, Thayer, Khalsa, & Lane, 2017; Stephan et al., 2016). This has considerable implications for mental health research, where disorders like schizophrenia, autism spectrum disorder or psychosomatic disorders have been linked to alterations in perception not only of the external world, but also the body (Adams, Stephan, Brown, Frith, & Friston, 2013; Friston et al., 2014; Haker, Schneebeli, & Stephan, 2016; Sahib S. Khalsa & Lapidus, 2016; Petzschner et al., 2017; Quattrocki & Friston, 2014; Stephan et al., 2016; Stephan & Mathys, 2014).

One framework that describes the intertwined nature of exteroceptive and interoceptive processes derives from the general notion of the “Bayesian brain”. Here, the brain is assumed to actively construct a generative model of its sensory inputs (from its external environment or from its own body), invert this model to determine the causes of its sensations (*inference),* continuously update the model (*learning*) based on new sensory information, and forecast future inputs (*prediction*). In other words, beliefs or predictions – probabilistic representations of environmental and/or bodily states – are updated based on sensory experience. Crucially, anatomically and mathematically concrete formulations of the different components of this framework exist (such as predictive coding, (Rao & Ballard, 1999) and active inference (Friston, 2009, 2010)), which suggest that learning or *belief updating* is prompted by mismatches between predictions and actual sensory inputs, formalized as prediction errors. Importantly, the weight that is given to any prediction error during the belief update depends on the relative precision assigned to the sensory channel (the low-level input) compared to the precision of (or confidence in) the (higher level) prior prediction. In other words, precise sensations increase and precise priors reduce belief updates.

Attention plays a prominent role in this framework. It has been conceptualized as a way to tune the relative weight of sensory information (prediction errors) on perceptual inference, both within and across different sensory modalities (Friston, 2009; Hohwy, 2012). More specifically, attention towards a specific sensory channel is thought to increase its relative precision and thereby the impact of the prediction errors it conveys (Feldman & Friston, 2010). Imbalances in this *precision weighting* (or salience assignment) have been proposed as key mechanisms in predictive coding accounts of psychiatric diseases (Adams et al., 2013; Friston et al., 2014; Haker et al., 2016; Lawson et al., 2014; Quattrocki & Friston, 2014; Stephan et al., 2016). Turning these ideas into clinically useful tests requires readouts of brain activity that reflect precision-weighted predictions errors both in the exteroceptive and the interoceptive domain. Such readouts must be robust and simple, and ideally generalize across cognitive contexts or task.

One candidate read-out in interoceptive processing is the heartbeat evoked potential (*HEP*), an electrophysiological brain response that reflects cortical processing of the heartbeat (Schandry, Sparrer, & Weitkunat, 1986). The HEP has previously been interpreted as an index of interoceptive belief updating; in particular, its trial-wise amplitude has been proposed to reflect a precision-weighted prediction error about each single heartbeat (Ainley, Apps, Fotopoulou, & Tsakiris, 2016).

If the HEP indeed represents as a neural correlate of interoceptive prediction error signals, then its amplitude should be modulated by attention. In particular, according to the predictive coding framework outlined above and analogous to previous studies on exteroception (Auksztulewicz & Friston, 2015), attention to interoceptive stimuli should heighten the precision of interosensory information and thus increase the weight of the associated prediction errors, relative to an attentional focus on exteroceptive channels. When attention is directed towards and away from the heart, this modulation should be reflected in the amplitude of the HEP.

In fact, most paradigms used in HEP research have implicitly probed some form of heart-related attention, for example, by asking participants to silently count heartbeats (*HbCounting task),* tap their fingers to each perceived heartbeat (*HbTapping task*), or discriminate between auditory stimuli presented in or out of synchrony with their heartbeat (*HbSync task*) (see Table 1 for an overview of these tasks and references to the literature). However, the modulation of attention in all of these tasks is assessed by posing additional task demands (e.g., counting or tapping) that may confound the interpretation of differences between interoceptive and exteroceptive attention. Examples include the additional auditory stimulation during the counting of tones compared to heartbeat counting, or the pronounced differences in difficulty (and thus performance levels) between conditions, given that exteroceptive stimulation is typically far above detection thresholds, which may lead to different task strategies (e.g., counting tones versus guessing heart rates).

**Table 1:** Overview of tasks used to probe cardiac perception

| Task | Description | Names in the literature | Example studies |
| --- | --- | --- | --- |
| **HbAttention** | P. focus their attention on their own heartbeat.<br>**Control condition:** P. focus their attention on a white noise sound (this paper), or a visual stimulus (Simmons et al., 2013) |  | Simmons et al. 2013**<br>This paper * |
| **HbCounting** | P. silently count their own heartbeats in different unknown time windows e.g. 25, 35 and 45 sec.<br>**Control condition:** P. count the number of (target) tones in different unknown time windows e.g. 25, 35 and 45 sec. | Heartbeat Perception Task, Heartbeat Tracking Task, Mental Tracking Task, Heartbeat Discrimination Task, Heartbeat Counting Task | Montoya, Schandry, Müller, et al. 1993*; Pollatos et al. 2005b*; Pollatos & Schandry 2004b*; Schandry, Sparrer, & Weitkunat 1986*; Terhaar et al. 2012* |
| **HbTapping** | P. tap a computer keyboard along with their perceived heartbeats. Often involves a pre-training, training (with feedback), and post training phase.<br>**Control condition:** P. tap a computer keyboard along with external stimuli (tones). Sometimes no control condition. | Heartbeat detection task | Canales-Johnson et al. 2015*; Garcia-Cordero et al. 2016*; García-Cordero et al. 2017*; Melloni et al. 2013; Sedeño et al. 2014**, Yoris et al. 2015, 2018* |
| **HbSync** | P. rate whether a sequence of stimuli (usually) tones is played in sync or out of sync / with a delay with respect to their heartbeat.<br>**Control condition:** P. detect target tones within the stream of stimuli (only in Critchley et al. 2004). Otherwise no control condition. | Heartbeat discrimination task, Heartbeat detection task | Brener & Kluvitse 1988; Critchley et al. 2004**; Garfinkel et al. 2015; Katkin et al. 1991*; Katkin, Reed, & Deero 1983; Sokol-Hessner et al. 2015; Wiens et al. 2000 |
| **HbDiscrimination** | P. determine whether an auditory recording was their own heartbeat or not. | - | Azevedo, Aglioti, & Lenggenhager 2016 |

*P. stands for participants. \* indicates EEG studies that measured an HEP. \*\* indicates fMRI studies*

Moreover, the vast majority of HEP studies do not report contrasts between exteroceptive and interoceptive conditions (Canales-Johnson et al., 2015; Garcia-Cordero et al., 2016; Katkin, Cestaro, & Weitkunat, 1991; Müller et al., 2015; Pollatos, Kirsch, & Schandry, 2005; Pollatos & Schandry, 2004; Schulz et al., 2015; Terhaar, Viola, Bär, & Debener, 2012; Werner, Jung, Duschek, & Schandry, 2009; Wiens, Mezzacappa, & Katkin, 2000). HEP differences between counting heartbeats and counting tones were presented by very few studies (Montoya, Schandry, & Müller, 1993; Schandry, Sparrer, & Weitkunat, 1986; but see Leopold & Schandry, 2001; Terhaar, Viola, Bär, & Debener, 2012 for nonsignificant results). One study reported a modulation of HEP amplitude by tapping to heartbeats compared to tapping to an external stimulus (García-Cordero et al., 2017). Surprisingly, to the best of our knowledge, there is not a single EEG study that examined changes in the HEP during a “pure” attention task contrasting interoceptive to exteroceptive attention without any additional task demands (in fMRI, Simmons et al., 2013, did use a blocked attention paradigm of this sort, but did not examine trial-wise heartbeat-related processing and also did not present the same exteroceptive stimulus during interoceptive processing). A clean demonstration of this modulation, however, is critical for justifying the use of the HEP as an interoceptive signal of precision-weighted prediction errors.

In this paper, we present an investigation of the HEP that uses two innovations to clarify this issue. First, we developed a novel heartbeat attention (*HbAttention*) task, which is designed to solely manipulate the attentional focus of participants (i.e., focusing either on their heart or on an external sound stimulus), without differences in stimulation and/or task. Second, given the considerable spatio-temporal variability of HEP effects in the literature – which have been reported between 171 ms (García-Cordero et al., 2017) and 595 ms (Schulz et al., 2013, 2015) after cardiac R peak events, and in frontal, central as well as parietal sensors – we performed an unbiased analysis covering the entire sensor space and the whole time window from 200 to 580 ms after cardiac R peak events while using a stringent correction for multiple comparisons.

## Methods

### Participants

Nineteen healthy male participants completed a newly developed Heartbeat Attention (*HbAttention*) task as part of a larger suite of other tasks which involved focusing on one’s own heartbeat or heartbeat-dependent stimulus presentation. The order of these tasks was counterbalanced across participants. All participants were right-handed, had normal or corrected-to-normal vision, and provided informed consent to participate in the study. An overview of the participants’ sociodemographic data is shown in Table 2. Furthermore, all participants fulfilled the following inclusion criteria: between 18 and 40 years old; no previous or current chronic disorder, injury, or operation related to the brain; including no history of neurological or psychiatric illness; no history of drug abuse; abstinence of medication or drug consumption seven days prior to the experiment; and abstinence of alcohol intake 24 hours prior to the experiment session. The protocol was approved by the Cantonal Ethics Committee Zurich (PB_2016-01717).

**Table 2:** Sociodemographic variables and questionnaire scores for the analyzed sample in this study.

|  |  |  |
| --- | --- | --- |
|  | **Mean (sd)** | **range** |
| Age | 23.68 (3.61) | 20 – 34 |
| Weight | 80.58 (12.17) | 65 – 116 |
| BMI | 23.95 (3.14) | 20.59 – 33.53 |
| Debriefing Questionnaire |  |  |
| Perceive Sound | 9.37 (1.01) | 7 – 10 |
| Perceive Heart | 5.11 (2.73) | 1 – 9 |
| Concentrate Sound | 9.11 (0.94) | 7 – 10 |
| Concentrate Heart | 6.37 (2.45) | 2 – 10 |
| BPQ |  |  |
| Body awareness | 1.85 (1.13) | 0.04 – 3.73 |
| Supradiaphragmatic reactivity | 0.68 (0.65) | 0 – 2 |
| Subdiaphragmatic reactivity | 0.29 (0.34) | 0 – 1.13 |
| **Notes: N=19** |  |  |

### Heartbeat attention (HbAttention) task

The HbAttention task consisted of alternating blocks of 20s, in which participants were instructed to focus their attention either on their own heartbeat (*interoceptive attention,* condition: *HEART*) or a sound stimulus (*exteroceptive attention,* condition: *SOUND).* Blocks were separated by a rating period (max duration: 9 s) and an inter-trial-interval (ITI) of varying length (5 s −15 s) drawn from a uniform distribution (Figure 1A).

**Figure 1:**
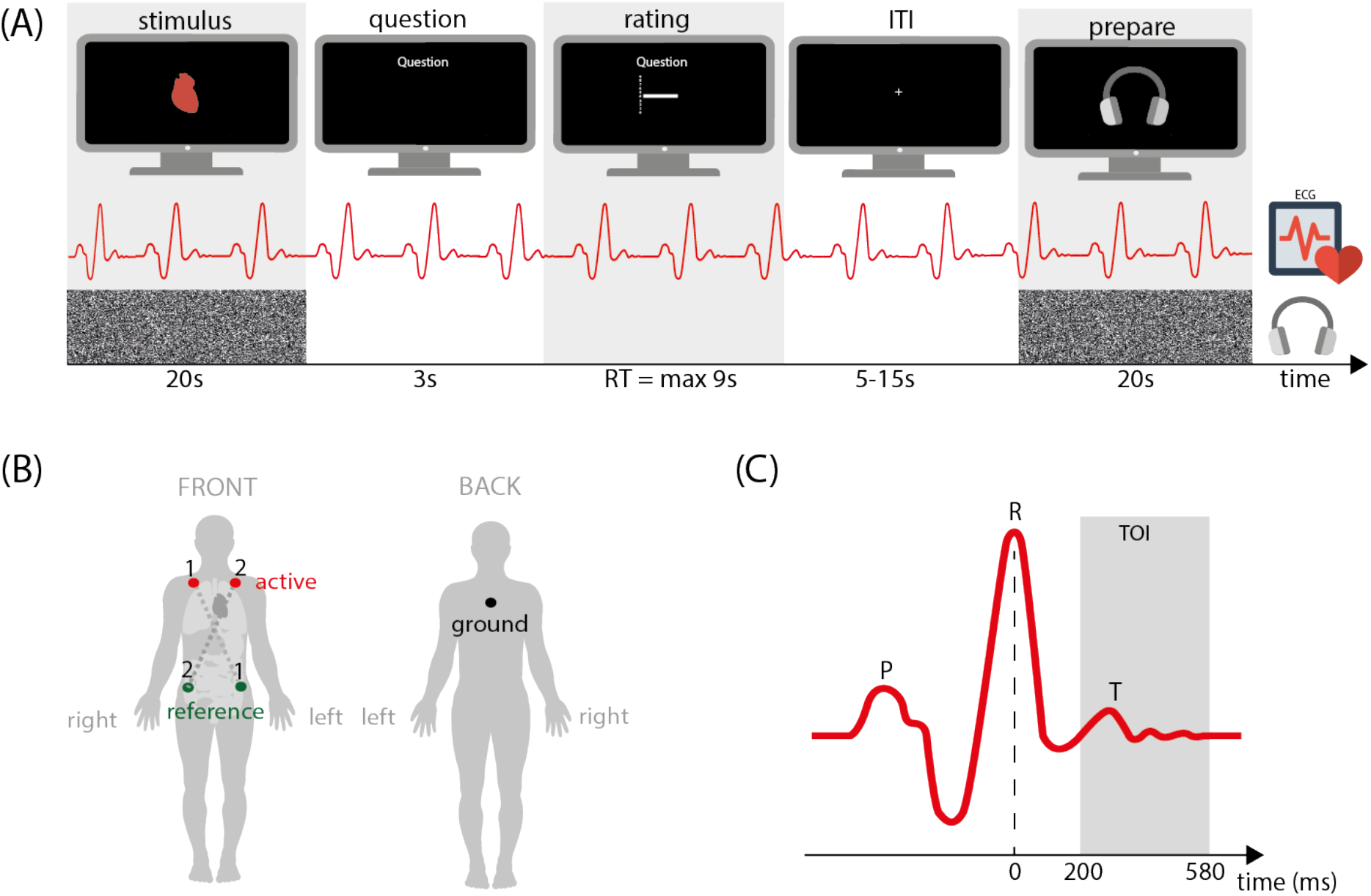
(A) Schematic overview of the block design in the heartbeat attention task (HbAttention). Blocks of attention to one’s own heart (interoceptive HEART condition) were alternated with blocks of attention to an auditory white noise stimulus, which was presented via head phones (exteroceptive SOUND condition). Conditions were separated by rating questions about the associations an individual had with the previous attention block, and an ITI of variable length. Importantly, both auditory sounds and heartbeats were present during both types of attention blocks; this prevented any sensory differences between conditions. (B) Placement of the electrodes in the two ECG derivations. ECG1 was used in all analyses for this paper. (C) Schematic pattern of ECG signal during a heartbeat, including the P wave, R peak and T wave. The gray area indicates the time window of interest (TOI) in which our statistical analyses of EEG data were conducted (200-580ms).

A HEART block was indicated by a visual heart symbol on screen (see Figure 1A), which was presented for the entire duration of the block. Participants were instructed to focus their attention on their heart as long as the heart symbol was displayed and to attend to changes in the sensation of the heart or their heart rate to the best of their abilities, without measuring their pulse. A SOUND block was indicated by a headphone symbol on screen, which was presented for the entire duration of the block. Participants were instructed to focus their attention on the white noise issued via headphones as long as a headphone symbol was displayed on screen, and to attend to any potential changes in the sound. Importantly, the white noise sound was played during all (HEART and SOUND) blocks to ensure that any observed changes in brain activity were consequences of a shift in attention and not due to changes in sensory stimulation.

The participants were also instructed that, in both conditions, they would be asked to rate specific aspects of their perception of or associations with the previous attention block. These rating questions varied across blocks and conditions (e.g., *‘How well were you able to concentrate on the white noise in the last block? ‘* or *,How much would you associate your perceived heart beat in the previous block with the color red?* ‘) and participants responded on a 1 to 10 scale. Notably, the only purpose of these questions was to keep participants alert and focused, providing an incentive for them to pay attention during the blocks. In an earlier pilot version of the task we did not include these questions and observed that the effect of attention was reduced. We did not include the ratings in any of the further analyses.

During the inter-trial-interval (ITI), which was indicated by a fixation cross, participants were free to think about whatever they wanted to. There was no sound stimulus during the ITI or the rating period. Each condition was repeated 10 times throughout the experiment in a pseudo-randomized order. The same order was kept constant across participants (H-S-H-S-S-H-H-S-H-S-H-S-S-H-S-H-H-S-H-S). The HbAttention task lasted about 12 min. The task was programmed in MATLAB (MathWorks, version: 8.5.1.281278 (R2015a)) and used Psychtoolbox (http://psychtoolbox.org/). The code for the HbAttention task will be made available as open source code in a future release of the TAPAS toolbox (www.translationalneuromodeling.org/software).

### Debriefing Questionnaire

Immediately after the HbAttention task all participants filled out a debriefing questionnaire (in German) to indicate how well they were able to perform the task. The debriefing contained four questions regarding how well participants could concentrate on and perceive their heart and the sound stimulus (Sound Perception: QSP = ‘Were you always able to perceive the sound (white noise on your headphones) well?’; Sound concentration QSC = ‘Were you always able to concentrate well on the sound during sound blocks?’; Heart perception: QHP = ‘Were you always able to perceive your heartbeat well?’; Heart concentration: QHC = ‘Were you always able to concentrate well on your heart during heart blocks?’) (see Table 2). Responses were given on a scale from 1 (‘not at all’) to 10 (‘very well’). In addition, participants were asked to indicate (yes/no) if they had used any aids to ‘feel their heartbeat better’ (e.g., by measuring their pulse). Three out of 18 participants confirmed this question. Two of these participants reported that they sometimes ‘closed their eyes to concentrate better on their pulse’ and one reported to have ‘concentrated on a body part where the pulse was felt more prominently’. None of the participants reported to have measured their pulse directly. Overall, the participants’ selfreported perception and concentration levels were in the mid-range for the HEART condition (Debriefing: mean heart perception score = 5.11 (*sd* = 2.73), mean heart concentration score = 6.37 (*sd* = 2.45)), and in the high-range for the SOUND condition on a scale from 1 to 10 (mean sound perception score = 9.37 (*sd* = 1.01), mean sound concentration score = 9.11 (*sd* = 0.94)) (see Table 2).

### Body Perception Questionnaire (BPQ)

Prior to the experimental session participants filled out a number of online questionnaires. For the heartbeat attention task reported in this paper, we examined the Body Perception Questionnaire (*BPQ,* (Porges, 1993)) to investigate whether attentional modulation as induced by our task would be predictive of self-reported measures of bodily awareness and reactivity.

The BPQ is a 122-item questionnaire relating to the polyvagal theory (Porges, 2007). It measures awareness and reactivity of the autonomic nervous system, i.e. the subjective ability to perceive bodily states and bodily reactions to stress. Additionally, the BPQ assesses socio-demographic data and substance use. High scores on the BPQ reflect high awareness of internal bodily signals (i.e., high interoceptive sensibility) and high perceived reactivity of the visceral nervous system. The BPQ can be analyzed regarding 3 sub-scales, namely body awareness, supradiaphragmatic reactivity, and subdiaphragmatic reactivity, where the latter two refer to questions about the reactivity of organs above and below the diaphragm, respectively (Cabrera et al., 2017). In this paper, we focus on the body awareness and supradiaphragmatic reactivity subscales, given our specific focus on attention to the heart.

### Data Acquisition

Stimuli were presented within an electromagnetically shielded, sound attenuated, dimly lit EEG cabin via a stimulus PC (Hardware: Axxiv SVELT AZ7701 MD, CPU: Intel Core i7 3770K, GPU: Nvidia GTX660 2 GB GDDR5, 1344 cuda cores, Audio: Asus Xonar Essence STX, OS: Windows XP SP3). Continuous EEG was recorded on a 64-channel BrainCap with multitrodes (EASYCAP GmbH, Herrsching, Germany) using DC-amplifiers (BrainProducts GmbH, Gilching, Germany). The reference electrode was placed on the tip of the nose. The positions of all 64 electrodes were digitized after the experiment was finished. One electro-oculogram (*EOG*) electrode was placed below the left external canthus to monitor vertical eye movements and blinks. In addition, two electrocardiograms (*ECG*) were acquired using two electrodes placed on the left and right clavicle (active electrodes), one electrode below the neck above the shoulder blades (ground electrode), and two electrodes placed at the left and right hip/abdominal (reference electrodes), respectively (Figure 1B). For a subset of participants, the second ECG was recorded at the inside of the arm just below the crook of the arm on both the left- and right-hand side. Both ECG derivations shared the same ground electrode but were otherwise analyzed and acquired independent of each other. The arrangement of the first ECG (right clavicle – left hip) was chosen to optimize the expression of both the cardiac R peak and T wave, to facilitate their detection online and offline. The second ECG (left clavicle – right hip) served as a back-up in case the first ECG signal quality would have been too low and R peak detection would have been unreliable. In the current dataset, however, the quality of the first ECG data was high for all participants, thus the data of the second ECG were not used in the analysis. Breathing was recorded using a respiration belt to measure the thoracic or abdominal movements (BrainProducts GmbH, Gilching, Germany) and skin conductance (GSR MR sensor, BrainProducts GmbH, Gilching, Germany) was recorded on the second segment of the left middle and index fingers. Both of these metrics were not analyzed in the context of this paper. All signals were recorded with a sampling rate of 500 Hz. The onset and offset of each block (HEART, SOUND, RATING and ITI) was marked by triggers in the recordings of EEG data.

### EEG and ECG Preprocessing

Data were analyzed using SPM 12 (r6906) and in-house software developed in MATLAB (MathWorks, version: 8.5.1.281278 (R2015a)). Sensor locations of all EEG channels were based on the custom template location of the caps used.

EEG data were filtered offline using a high-pass filter (zero-phase shift two-pass Butterworth filter with cutoff at 0.5 Hz), then downsampled to 250 Hz and filtered again with a low-pass filter (zero-phase shift two-pass Butterworth filter with cutoff at 30 Hz). ECG data were downsampled to 250 Hz. Cardiac R peak events were detected for every heartbeat in the downsampled ECG using an online sample software package (http://www.librow.com/articles/article-13) based on a fast Fourier transform combined with an in-house extension. R peak times were saved separately for HEART and SOUND blocks for subsequent analyses (see section “Tests to exclude confounding cardiac effects” below).

EEG data were epoched with the R peak event as temporal reference (epoch length: −100 to 580 ms after R peak). As mentioned in the introduction, previous HEP reports showed a considerable variability with respect to the timing and topography of the HEP. We therefore analyzed an extended time window of interest (TOI) across the entire sensor space and applied multiple comparison correction, as described below. The TOI for the statistical analysis was set to 200-580 ms (Figure 1C). The choice of onset for the TOI was guided by three reasons: it approximately corresponds to (i) the earliest time point at which HEP effects have previously been reported (García-Cordero et al., 2017), (ii) the earliest time point at which participants in previous studies reported (consciously) sensing external stimuli as being synchronized to their heart (Brener & Ring, 1995; Wiens et al., 2000), and (iii) the time of increased baroreceptor firing when systolic blood outflow stretches the wall of the aortic arch and carotid sinus (Gray, Minati, Paoletti, & Critchley, 2010). The chosen end point of the TOI corresponds approximately to the latest time point when HEP components have been reported previously (Schulz et al., 2013, 2015) and was picked to assure that the TOI did not overlap with early components of the cardiac field artefact (*CFA*) of the next heartbeat, which originates from the electric field caused by the contraction of the heart muscles, for most heartbeats (trials in which this was the case were later excluded from the analysis). For the same reason we also did not baseline correct our epochs as any chosen baseline period would very likely have been confounded by components of the CFA. In particular, the P and Q waves, which occur just before the R peak, would be present in typically chosen baseline periods (Figure 1C). In addition, in periods of high heart rates (small R-to-R intervals) the time window right before the R peak, which is usually used for baseline correction, could potentially overlap with late components of the HEP, which have been reported up to 595 ms after the R peak (Schulz et al., 2013, 2015).

We detected eye blinks by thresholding the detrended and filtered EOG channel as implemented in SPM12. Epochs in which the heartbeat (within a window of 100-600 ms after R peak) overlapped with an eye blink (assuming an average eye blink duration of 500 ms) were rejected. On average, 61 (*sd* = 53) epochs were rejected due to eye blinks, of which an average of 24 (*sd* = 23) epochs were part of the HEART condition and 38 (*sd* = 32) epochs were part of the SOUND condition. There was no significant difference in the number of rejected epochs due to eye blinks in the SOUND condition and HEART condition (Wilcoxon Signed-Ranks Test: *Z* = −1.90, *p* = 0.06). In addition, to avoid a contamination of our TOI (200-580 ms) by the CFA of the P wave or QRS complex of the following heartbeat in periods of high heart rates, we rejected all trials with an R-to-R interval below 630 ms. Epochs associated with short R-to-R intervals were only observed for three out of nineteen participants, and for these three, we had to reject 2, 13 and 40 epochs, respectively. Finally, we rejected all epochs in which the signal recorded at any channel exceeded a threshold of +/-75 μV and marked channels as bad in which the proportion of rejected trials exceeded 20%. On average, 14 epochs (*sd* = 16) were rejected due to exceedance of the amplitude threshold. No channels were marked as bad.

The CFA represents an important potential confound for investigations of the HEP, with no universally agreed upon solution. One generally powerful method for artefact correction of EEG data is independent component analysis (ICA) (Debener, Thorne, Schneider, & Viola, 2010; Devuyst, Dutoit, Stenuit, Kerkhofs, & Stanus, 2008; Hoffmann & Falkenstein, 2008). This method has its caveats and we are not aware of any study that demonstrated a full removal of the CFA by means of ICA: EEG data after ICA correction often show a remaining R peak of varying size caused by the heart’s electrical field; in addition, the T wave artefact, which is prominent in many channels and overlaps in time with the early part of the TOI, is rarely completely corrected by ICA. Additionally, ICA can cause a removal of task-relevant signals. Fortunately, in our particular paradigm, we contrast the HEP across two attentional conditions. In the absence of any differences in cardiac activity between conditions (see below), it is safe to assume that the CFA is constant across the two conditions of our task and will therefore not affect the contrast between HEART and SOUND conditions. We conducted several analyses to test this assumption, by comparing the heart rate and ECG signal amplitude across conditions in addition to the classical EEG analysis (see below: Tests to exclude confounding cardiac effect).

After artefact rejection, we included, on average, 371 epochs (*sd* = 68) per participant in the ERP analysis. The number of included epochs did not differ significantly between the HEART (*mean* = 191, *sd* = 32) and SOUND (*mean* = 180, *sd* = 37) condition (Wilcoxon Signed Rank Test: *Z* = 0.91, *p* = 0.37).

### Group-level EEG analysis

For statistical analysis, all included epochs were averaged within participants separately for both conditions and converted to 2D (32 × 32 pixel) scalp images for all 96 time points (under the sampling frequency of 250 Hz) within the window of interest. This resulted in a 3D image per condition and participant, using a voxel size of 4.2 mm × 5.4 mm × 4.0 ms. The 3D space-time images were spatially smoothed using a Gaussian kernel (FWHM: 16 mm × 16 mm) in accordance with the assumptions of Random Field Theory (Kiebel & Friston, 2004; Worsley et al., 1996) to accommodate for between-subject variability in channel space and entered into a General Linear Model (GLM) on the group-level.

Differences in HEP amplitude between the two conditions (HEART vs. SOUND) were examined using a paired t-test. We report any effects that survived whole-volume (sensor × time space) family-wise error (FWE) correction at the cluster-level (p < 0.05), with a cluster-defining threshold of p < 0.001.

### HEP amplitude analysis

In addition, we calculated the average HEP amplitude (*HEPa*) per condition and participant. While in classical ERP analyses this is usually done by averaging the signal of the most significant electrode(s) in a specific time window, here we chose a less restricted approach and included all significant parts of the sensor × time space that survived multiple comparison correction in the SPM analysis. To that end, we computed a 3D (sensor × time) mask that included all significant voxels from the contrast HEART versus SOUND in the group-level EEG analysis. Using this mask, we selected each voxel’s activity from the smoothed 3D images for each participant and each condition and then calculated an individual’s average HEPa for the HEART and SOUND condition.

To relate inter-individual differences in the effect of attention on the HEPa to external questionnaire-based measures, we calculated the participant-specific HEPa difference (ΔHEPa) between the conditions, by subtracting the average HEPa during SOUND blocks from the average HEPa during HEART blocks. Finally, we ran a linear regression analysis, predicting individual’s scores on two subscales of the BPQ (body awareness and supradiaphragmatic reactivity) using the ΔHEPa.

### Tests to exclude confounding cardiac effects

To verify our assumption that the CFA would equally impact the HEP in both conditions, and that any difference in EEG amplitude could therefore not be attributed to differences in the electric field of the heart itself, we calculated the individual average ECG amplitude (ECGa) within the time window of significant HEP differences for each condition and tested for a difference between HEART and SOUND conditions using a Wilcoxon Signed-Rank Test. In addition, we tested if there was any relationship between the ECGa and HEPa differences across participants using a linear regression to predict the ΔHEPa from the ΔECGa. Finally, we also tested for a significant difference in heart rate (ΔHR) across the two conditions using a Wilcoxon Signed-Rank Test.

### Statistical tests for non-EEG variables

Statistical analyses were performed in MATLAB (MathWorks, version: 8.5.1.281278 (R2015a)). For comparisons across experimental conditions (HEART versus SOUND) outside the SPM-based EEG analysis we first used a single sample Kolmogorov-Smirnov test for all variables of interest to determine whether they were normally distributed (Matlab function *kstest).* In case they were not, we ran a non-parametric Wilcoxon Signed Rank Test using (Matlab function *<u>ranksum</u>*). Linear regression models were calculated using Matlabs’s *fitglm.* A probability level of*p* < 0.05 was considered significant for all statistical analyses. Bonferroni corrected p-values were reported for the regression analysis between the ΔHEPa and the two analyzed subscales of the BPQ (body awareness and supradiaphragmatic reactivity).

## Results

Using an unrestricted approach that included the entire sensor × time space plus correction for multiple comparisons, we tested for changes in the HEP as a function of the focus of attention. We found a significant difference in the HEP amplitude between attention to the interoceptive versus exteroceptive stimulus: The HEP amplitude was significantly higher during interoceptive compared to exteroceptive attention in a time window of 520 to 580 ms (peak at 576 ms after R peak) over central right channels (centered around C4 and CP4, T(1,18): 4.92; *p* < 0.05 corrected, see Table 3, Figure 2 and Figure 3A,B).

**Figure 2:**
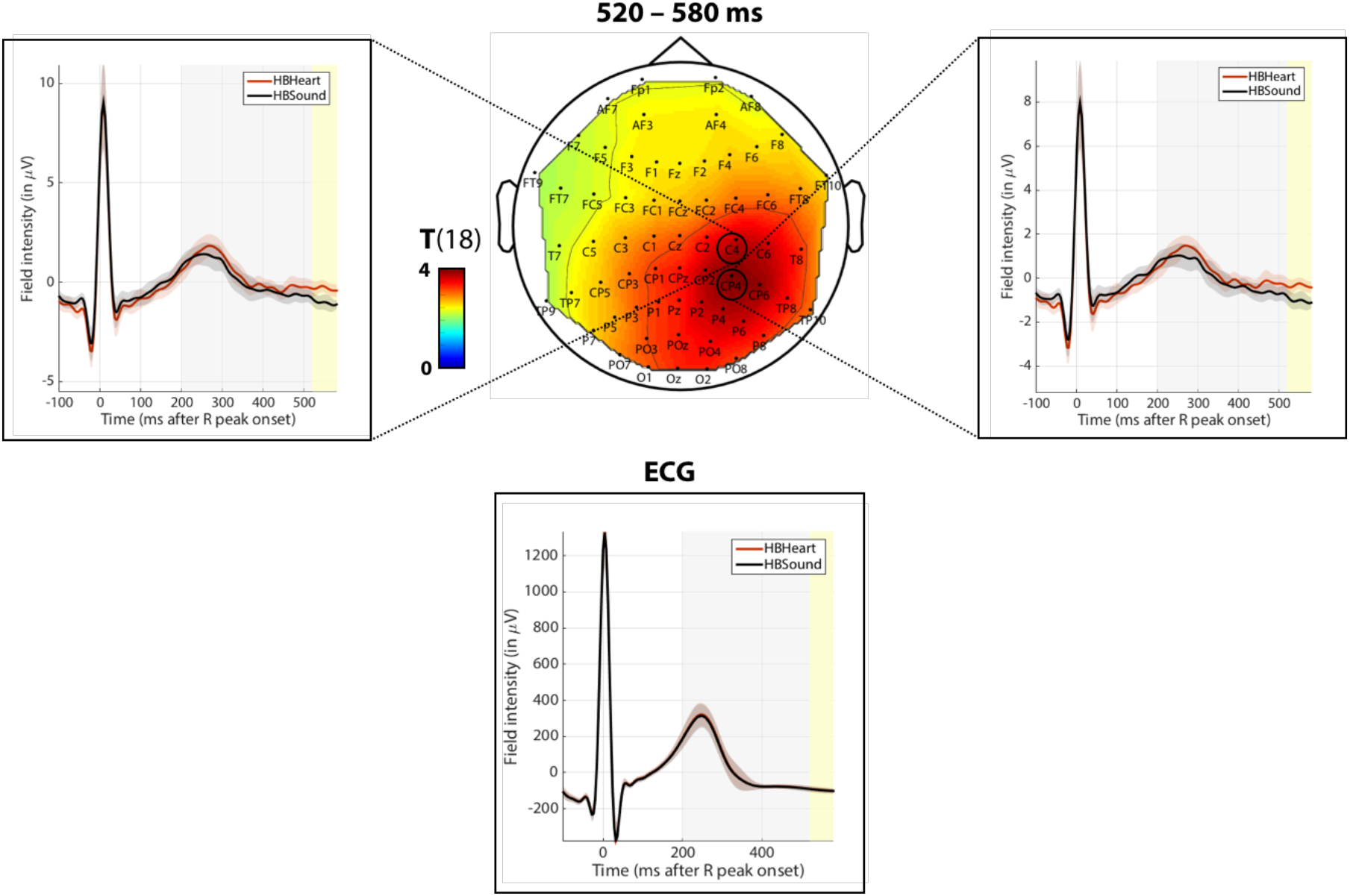
Middle: Scalp map of the average T-statistic for the contrast HEART >SOUND in the full statistically significant time window of520-580ms. Left and Right: Example ERP waveform for an electrode **within** the significant cluster of activation (Left: C4, Right: CP4). Bottom: Averaged, epoched ECG signal in the same time window. The HEART condition is indicated in red, the SOUND condition in black. The gray rectangle indicates the temporal window of interest used for the statistical analysis (TOI: 200 - 580 ms after R peak), the yellow rectangle marks the time window of significant activation (520 – 580 ms after R peak). Time is indicated with respect to the timing of the R peak. Shaded error bars indicate 95% confidence intervals.

**Figure 3:**
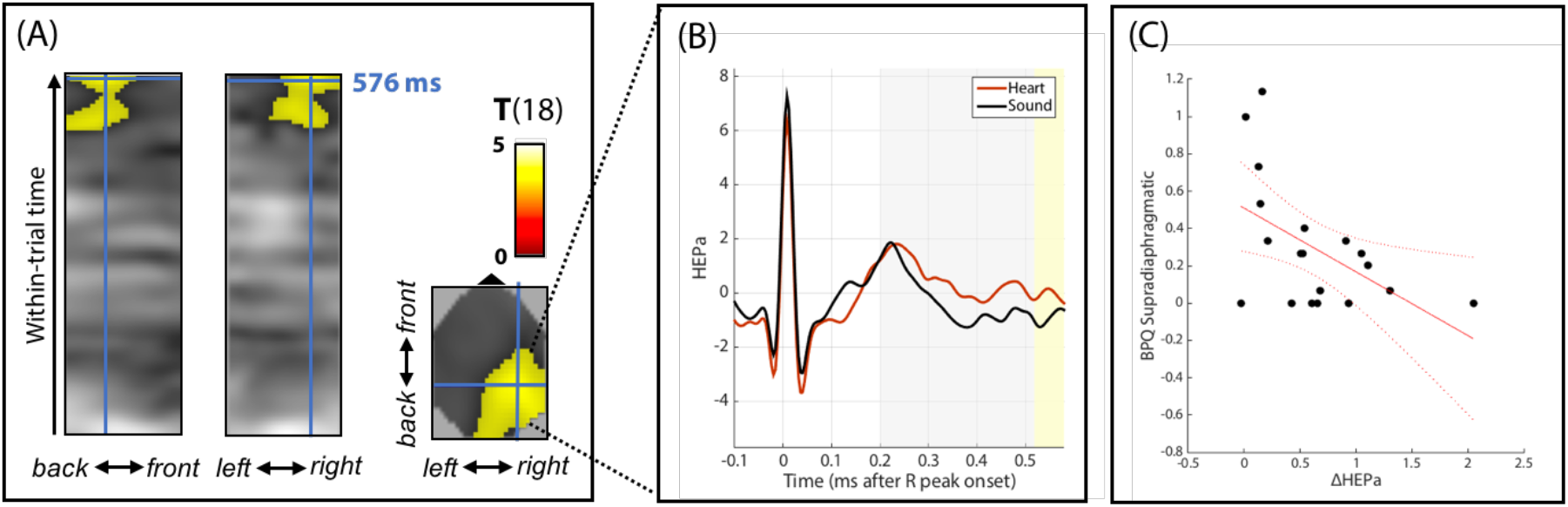
(A) Space x time statistical parametric maps. Yellow mask depicts significant activation based on the t-test for HEART >SOUND (whole brain FWE corrected at p < 0.05 at the cluster-level, with a cluster-defining threshold of p<0.001). The blue crosshair indicates the time and location of peak activation at 576ms. (B) Average HEP amplitude for the HEART (red) and SOUND (black) condition over time in an x-y mask including all significant voxels in sensor space over time. The plot was done by averaging these voxels ‘ activity from the smoothed 3D images for every participant, condition and time point. The gray rectangle indicates the window usedfor the statistical analysis (TOI, 200-580 ms after R peak), the yellow rectangle marks the time window of significant activation (520-580 ms after R peak). (C) Scatter plot of the linear relationship between the average HEP height and the supradiaphragmatic subscale of the BPQ. The red line indicates the linear fit including the 95% confidence bounds (dotted lines).

**Table 3:** Test statistics for the effect of attention in the contrast HEART > SOUND on EEG amplitude

| *Activation size*<br>(voxels) | *Cluster*<br>*p-value*<br>(FWEcorr) | *Peak*<br>*p-value*<br>(FWEcorr) | *Peak*<br>*T statistic* | *Peak*<br>*Z statistic* | *Peak coordinates*<br>(in mm) | *Peak latency*<br>(ms) | *Sign. time*<br>*window*<br>(ms) |
| --- | --- | --- | --- | --- | --- | --- | --- |
| 1988 | 0.009* | 0.049* | 4.92 | 3.86 | x = 34 , y = -36 | 576 | 520 - 580 |

Significant FWE-corrected *p*-values are indicted by an asterisk. Latencies are provided with respect to the R peak onset.

Although our design (and the late temporal expression of the observed difference) makes a confounding influence of CFA unlikely (see Methods), we performed additional analyses to exclude that the observed difference in HEP amplitude between interoceptive and exteroceptive attention could have been driven by physiological (cardiac) differences between conditions. First, we tested for differences in ECG amplitude and heart rate, respectively, across the two conditions. We found no significant effect of condition on the ECG amplitude in the time window of significant EEG effects (Wilcoxon Signed Rank Test on ΔECGa: *Z* = 0.06, *p* = 0.95), nor on the heart rate (Wilcoxon Signed Rank Test: *Z* = −0.61, *p* = 0.54) (Figure 4A, B). This suggests that the observed attentional effects on HEP amplitude could not be explained by physiological changes of heart function across conditions.

**Figure 4:**
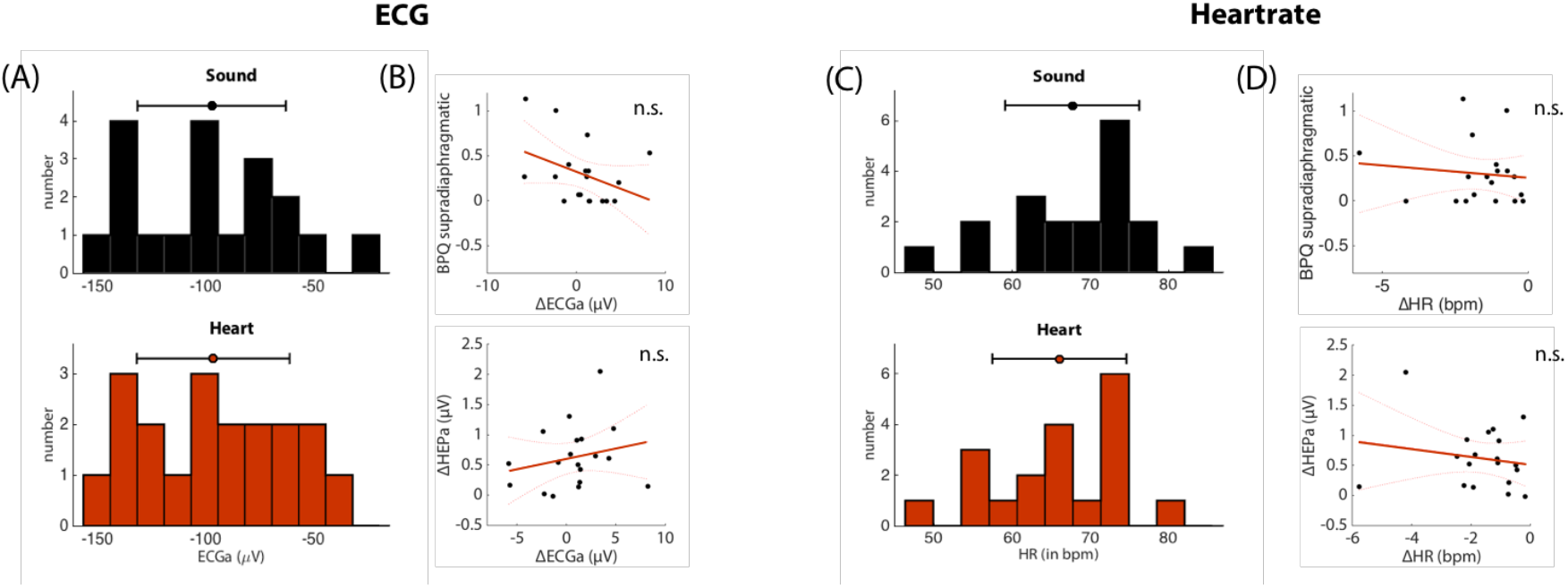
Sub-set of additional tests to exclude that the results are driven by cardiac confounds. (A) and (C): Histogram of the ECG amplitude (ECGa) in the significant time window (520-580ms after R) andheartrate (HR) across the conditions SOUND (black) and HEART (red). Top bar indicates mean and confidence interval. There was no significant difference for both ECG amplitude or HR between the attention to interoceptive and exteroceptive stimuli. (B) & (D): Scatter plot of HR difference and difference in ECG amplitude (ΔECGa) (HEART - SOUND) with BPQ supradiaphragmatic reactivity (upper plots) and the difference in HEP amplitude (ΔHEPa) during the significant time window (lower plots). The red line indicates the linear fit including the 95% confidence bounds (dotted lines). Individual differences in HEP amplitude or BPQ supradiaphragmatic reactivity could not be explained by individual differences in ECG amplitude or heartrate (see text for results of the statistical analysis).

Next, we tested if individual differences in HEP amplitude were related to differences in self-reported perception or concentration levels during the task. We found that even though self-reported concentration levels and perception in the debriefing differed across conditions (Wilcoxon Signed Rank Test Perception: *Z* = −4.67*,p* < 0.001; Wilcoxon Signed Rank Test Concentration: *Z* = −3.66*,p* < 0.001), they were not associated with differences in HEP (linear regression; Concentration: *F* = 3.46, *p* = 0.08; Perception: *F* = 0.78, *p* = 0.39).

Moreover, we tested if individual differences in HEP amplitude were associated with external questionnaire-based measures of interoception. To that end, we used a linear regression model with individual HEP amplitude differences to explain participants’ scores on the body awareness and supradiaphragmatic reactivity subscale of the BPQ. We found a significant negative relationship between the attention-induced difference in HEP amplitude and the supradiaphragmatic reactivity subscale of the BPQ (linear regression: *F* = 6.21, *p* = 0.02, *p_Bo_nf_e_rronì* < 0.05) (Figure 3C), such that higher HEP amplitude differences were associated with lower BPQ scores. By contrast, we found no significant relationship with the body awareness subscale of the BPQ (linear regression: *F* = 0.02, *p* = 0.88).

Finally, we revisited the possibility that the observed attentional effects may have been driven by individual variations in cardiac physiology. Specifically, we tested if individual differences in HEP amplitudes could be explained by individual differences in ECG amplitudes. We found no significant relation of this sort (linear regression: *F* = 0.89, *p* = 0.36). In addition, individual scores on the supradiaphragmatic reactivity subscale of the BPQ could not be explained by the average heart rate of an individual either (F = 1.2, *p* = 0.30), nor by the differences in heart rate across conditions (F = 0.23, *p* = 0.63), the average ECG amplitudes in the significant time window (F = 1.2, *p* = 0.28), or ECG amplitude differences (F = 2.8, *p* = 0.11) (Figure 4).

## Discussion

Using a new heartbeat attention (HbAttention) task designed to manipulate the attentional focus of participants without potentially confounding task demands, we found an increase in the amplitude of the HEP in a late time window (520-580 ms after R peak) during interoceptive compared to exteroceptive attention. Given the lack of differences in heart rate and ECG amplitude between conditions and the late timing of our effect, it is unlikely that this increase could have been driven by changes in cardiac activity and instead reflects a veridical effect of attentional focus. This was further corroborated by analyses that took into account independent measures of interoception: HEP amplitude differences between interoceptive and exteroceptive attention were significantly associated with self-reported supradiaphragmetic reactivity as measured by the BPQ, such that stronger attentional modulation of HEP amplitude was associated with smaller supradiaphragmetic BPQ scores. By contrast, we failed to find any relation between cardiac activity itself (as measured by ECG) and the supradiaphragmetic BPQ score (see Figure 4).

While attentional effects on the HEP have been reported previously (García-Cordero et al., 2017; Montoya, Schandry, & Müller, 1993; Schandry, Sparrer, & Weitkunat, 1986), this study is novel in two ways. First, it directly contrasts interoceptive to exteroceptive attention without preselecting a particular group of participants. Typically, HEP amplitudes have been contrasted between good and poor heartbeat perceivers, as defined by their performance on heartbeat detection tasks (HbCounting or HbSync, see description in Table 1) (Katkin, Cestaro, & Weitkunat, 1991b; Montoya, Schandry, & Müller, 1993; Pollatos, Kirsch, & Schandry, 2005; Pollatos & Schandry, 2004; Schandry, Sparrer, & Weitkunat, 1986). Most of these studies found that higher interoceptive accuracy (i.e. ‘good heartbeat perception’) was linked to higher HEP amplitudes during interoceptive conditions (Pollatos et al., 2005; Pollatos & Schandry, 2004; Schandry, Sparrer, & Weitkunat, 1986). However, these findings need to be interpreted carefully.

In addition to concerns about the interpretability of performance scores from existing heartbeat detection tasks (Brener & Ring, 2016; Ring & Brener, 2018), contrasting groups with differential heartbeat detection abilities does not allow for interpreting the observed effects as specific to the dynamics of internally-driven processes, unless there exists a specific comparison with exteroceptive conditions (García-Cordero et al., 2017).

Secondly, the relatively few studies that did directly contrast exteroceptive to interoceptive attention employed an additional task, typically heartbeat counting, which may have confounded the attentional effects. Montoya et al., for instance, used the HbCounting task and demonstrated differences in HEP amplitude between counting heartbeats and counting auditory events in a time window of 450 to 550 ms after the cardiac R peak (Montoya, Schandry, & Müller, 1993). However, the two conditions also differed with respect to the auditory stimulation, i.e., while heartbeats are always present, the auditory stimulus was only present during the exteroceptive condition. Furthermore, the conditions also varied with respect to their difficulty (i.e., usually it is easier to report tones than heart beats). This means that the observed HEP differences might not necessarily reflect a difference in attentional focus, but could be driven by the fact that participants only engaged in actual counting during the exteroceptive condition, for example, because they found it hard to consciously perceive any heartbeats. In the latter case, counting performance in the HbCounting task would either reflect the accuracy of the estimate of their own average heart rate, or simply their time estimation ability, rather than a beat-to-beat detection ability; this possibility has been pointed out previously (Ring & Brener, 2018; Ring, Brener, Knapp, & Mailloux, 2015).

García-Cordero and colleagues used a heartbeat tapping (HbTapping, see description in Table 1) task and showed HEP differences at numerous time points between 200 and 500 ms after the cardiac R peak between interoceptive conditions (before and after a veridical feedback session) and exteroceptive conditions where participants were instructed to tap to a simulated heartbeat sound (García-Cordero et al., 2017). The HbTapping task offers a richer readout in terms of heartbeat detection performance scores than the HbCounting task as it measures the temporal relation of every actual and perceived beat. However, it is confounded by the motor activity associated with tapping. Even though participants in García-Cordero et al. had to tap in both conditions, the clear difference in performance accuracy between exteroceptive and interoceptive conditions likely led to a differential effect of the motor artefact on EEG signal during epochs of interest, which hampers the interpretation of any EEG difference as a pure attention effect.

There are only very few studies that have examined the contrast between exteroceptive and interoceptive attention in a setting where only the focus of attention was manipulated. To our knowledge, none of these studies uses EEG or adopts a single-trial perspective on perception of cardiac activity. For example, fMRI studies found an activation for ‘pure’ interoceptive attention in subregions of the insular cortex, an area thought to represent the primary locus for interoceptive signal processing and multisensory integration of interoceptive and exteroceptive information (Farb, Segal, & Anderson, 2013; Simmons et al., 2013). Interestingly, the insula has also been previously localized as a potential source for the HEP (Park, Correia, Ducorps, & Tallon-Baudry, 2014; Park et al., 2017; Pollatos et al., 2005). The pure attentional effect observed in the current study is not confounded by differences in auditory or interoceptive stimulation itself (as the sound stimulus are present during all conditions, as is the heartbeat, naturally), our paradigm did not include any potentially confounding task demands or motor responses, and we observed no differences in cardiac activity across conditions.

Why is it relevant to study the relationship between the HEP and pure interoceptive attention? As mentioned in the introduction, the HEP is an interesting candidate for empirically testing predictions made by recent theoretical frameworks that understand perception of body and world in a joint Bayesian framework, in particular predictive coding (Rao & Ballard, 1999) and active inference (Friston, 2009, 2010). In the exteroceptive domain, a large body of psychophysical experiments has provided clear evidence for Bayesian inference during basic perceptual judgements and multi-sensory integration (for overviews see (Geisler & Kersten, 2002; Knill & Richards, 1996; Petzschner, Glasauer, & Stephan, 2015). Among others, recent studies in vision (Kok & De Lange, 2014; Muckli et al., 2015; Pinto, van Gaal, de Lange, Lamme, & Seth, 2015) and audition (Chennu et al., 2013) demonstrate how prior expectations shape behavioral and neuronal signatures of perception (De Lange, Heilbron, & Kok, 2018). Model-based analyses of behavioral and neuroimaging data provide evidence for a key feature of Bayesian belief updating (for distributions from the exponential family), i.e., precision-weighted prediction errors in various (exteroceptive) contexts (Diaconescu et al., 2014, 2017; Iglesias et al., 2013; Sedley et al., 2016; Stefanics, Heinzle, Horváth, & Stephan, 2018), suggesting that precision-weighting is a generic computational process throughout the brain. Finally, several studies support an interpretation of attention as an optimization of this precision-weighting (Jiang, Summerfield, & Egner, 2013; Vossel, Bauer, et al., 2014; Vossel, Mathys, et al., 2014).

However, while proposals that interoception follows the same Bayesian inference mechanisms are accumulating (for reviews, see (Barrett & Simmons, 2015; Chanes & Barrett, 2016; Critchley & Garfinkel, 2017; Gallagher & Allen, 2018; Gu, Hof, Friston, & Fan, 2013; Owens, Allen, Ondobaka, & Friston, 2018; Seth, 2013; Seth & Friston, 2016)), strong empirical evidence for interoceptive predictions or prediction errors is still lacking. According to predictive coding accounts of interoception, neural correlates of interoceptive processing should be modulated by attention, such that interoceptive attention heightens the relative precision of bodily sensory information (and thus increases the weight of the associated precision-weighted prediction errors) relative to exteroceptive attention, where the salience of bodily sensory information is downregulated. Our demonstration of a purely attentional modulation of the HEP is thus consistent with its interpretation as a neural signature of interoceptive prediction errors (Ainley et al., 2016).

Moreover, using the individual strength of this modulation – the difference in HEP amplitude between interoceptive and exteroceptive conditions – we were able to explain a significant amount of variance in self-report measures (BPQ) of supradiaphragmatic autonomous nervous system reactivity (Figure 3C). Again, this relation was not due to individual variation in cardiac activity itself (as measured by the ECG amplitude, heart rate, or their changes across conditions) (Figure 4). High scores on this BPQ subscale indicate high reactivity of supradiaphragmatic organs such as the heart, related to the perceived frequency of potential warning or illness signs from these organs (e.g., *‘My heart is beating irregularly ‘, ‘I feel short of breath’).* In our sample, high scores on this scale were associated with small differences in the HEP between interoceptive and exteroceptive conditions. One possible explanation of this finding would be that a heightened (or more frequent) perception of interoceptive signals is caused by an inability to downregulate the precision (or salience) of these signals in situations where attention towards the body is not required. This inability to optimize the precision weights of different sensory channels during different task demands seems to be reflected in a reduced attentional modulation of the HEP in our task, lending further support to its potential utility as an individual readout of interoceptive processing.

One caveat for the interpretation of this result in the regression analysis is our relatively small sample size. The reliability of the ERP results, however, profits from the large number of trials per participant (even after exclusion of short R-to-R trials, eye blink trials, and other artefacts as detected by amplitude threshold).

Nevertheless, further evidence for the validity of the HEP as a quantitative metric for interoceptive information processing and, in particular, prediction error signaling, will be needed. These tests may require targeted manipulations to introduce interoceptive surprise about the heart or other internal organs, e.g. via the stimulation of vagal or other autonomic nerves (Borovikova et al., 2000; Burger et al., 2017; Nonis et al., 2017), by altering cardiac activity pharmacologically, using sympathomimetics with short half-life such as Isoprotenerol (Hassanpour et al., 2016), or by manipulating breathing perception by changing breathing resistance (Faull, Hayen, & Pattinson, 2017; Faull & Pattinson, 2017; Vinckier, Morélot-Panzini, & Similowski, 2018). An alternative strategy involves developing more sophisticated experimental designs and analysis strategies that allow us to probe (the variability in) brain responses to internal signals on a trial-by-trial basis or even induce interoceptive prediction errors via carefully designed exteroceptive stimulation (Marshall, Gentsch, Jelinčić, & Schütz-Bosbach, 2017; Owens et al., 2018; Van Elk, Lenggenhager, Heydrich, & Blanke, 2014). Finally, high resolution fMRI could become an additional important tool to resolve layer-specific hierarchical message passing of predictions and prediction errors in the cortex (Heinzle, Koopmans, den Ouden, Raman, & Stephan, 2016; Kok, Bains, Van Mourik, Norris, & De Lange, 2016; K.E. Stephan et al., 2017).

Compared to the exteroceptive domain, the extension of ‘Bayesian brain’ ideas to the interoceptive domain is more recent (Ainley et al., 2016; Barrett & Simmons, 2015; Ondobaka, Kilner, & Friston, 2017; Petzschner et al., 2017; Quattrocki & Friston, 2014; Seth & Critchley, 2013; Seth & Friston, 2016; Seth et al., 2012; Stephan et al., 2016). This framework has inspired a reconceptualization of complex aspects of “being human”, such as the neuronal basis of emotions (Gallagher & Allen, 2018; Seth, 2013; Seth & Friston, 2016) or selfhood (Seth, 2013; Seth et al., 2012). Equally importantly, this idea has led to new formalizations of clinically highly relevant concepts, such as allostatic control (Stephan et al., 2016). More generally, it is of considerable clinical relevance for understanding the origin and nature of psychosomatic symptoms, as highlighted in several recent articles (Khalsa et al., 2018; Khalsa & Lapidus, 2016; Owens et al., 2018; Paulus & Stein, 2010; Petzschner et al., 2017; Quattrocki & Friston, 2014; Sedeño et al., 2014; Klaas E Stephan et al., 2016).

In practice, quantitative read-outs of interoceptive information processing could become valuable diagnostic tools for detecting aberrant attention to interosensations and ensuing changes in interoceptive prediction errors. For example, overly salient sensory signals from the body have been proposed for certain types of anxiety, panic disorder and hypochondriasis, while suppression of bodily inputs (e.g., due to dominating effects of predictions) have been suggested for certain types of depression (Domschke, Stevens, Pfleiderer, & Gerlach, 2010; Dunn et al., 2010; Dunn, Dalgleish, Ogilvie, & Lawrence, 2007; Paulus & Stein, 2010; Stephan et al., 2016). In fact, alterations of the HEP have already been reported in a number of psychiatric and neurological conditions (García-Cordero et al., 2017; Müller et al., 2015; Schulz et al., 2015; Terhaar et al., 2012). If the findings from the present study – that the HEP is modulated by pure attention – and the interpretation of the HEP as a neural correlate of interoceptive prediction error signaling can be corroborated by future work, HEP recordings (and model-based analyses thereof) may develop into clinically useful tools for psychosomatics.

## Acknowledgements

We gratefully acknowledge generous support by the René and Susanne Braginsky Foundation and the University of Zurich.

